# Public engagement with biomedical science: An analysis of Black, Hispanic, and General Population adults in the United States

**DOI:** 10.1101/2023.03.15.530692

**Authors:** Adina T. Abeles, Kishore Hari, Cory Manento, Kyle Block, Stefan Musch, Jonathon P. Schuldt, Tania Simoncelli

**Affiliations:** Chan Zuckerberg Initiative; Gradient Metrics, LLC; Cornell University

## Abstract

The COVID-19 pandemic has highlighted the need to better understand public engagement with biomedical science across groups–especially those that have been historically marginalized by the medical science community. However, common approaches to studying group differences in science attitudes are often limited by small sample sizes or by categorizing individuals based on demographic variables, which may obscure meaningful variability within a given population. We recruited three probability-based samples of Black (N = 963), General Population (N = 957), and Hispanic (N = 964) adults living in the U.S. (October 21 - November 5, 2021). Employing a novel application of a non-negative matrix factorization model to conduct an attitudinal-based segmentation that categorized individuals based on distinct orientations toward biomedical science, our analysis revealed 10 unique mindsets across the three surveyed populations. Overall, our work underscores the value of recruiting independent samples from underserved and marginalized communities by revealing underexplored variation in how different publics orient toward biomedical science.

**Teaser:** Using a values-based segmentation approach, we uncover 10 unique biomedical mindsets in the general, Black, and Hispanic adult populations in the U.S.

## Introduction

Since the early days of the COVID-19 outbreak, understanding the factors that shape the public’s attitudes toward biomedical science has been widely regarded as a linchpin for the success of public health interventions and long-term engagement in science, as evidenced by the numerous studies examining the role of trust (and mistrust) in science. (e.g., 1-3;). Given the global nature of the pandemic, some studies have taken a cross-national approach to analyzing how citizens orient toward science. For instance, in a study involving nearly 7,000 participants from 23 countries, trust in science emerged as a key predictor of intentions to enact behaviors that protected oneself and one’s community from the coronavirus (4). At the same time, there are reasons to study engagement with science within particular countries, such as the United States, as we do in the present research. Different cultural norms, laws, scientific and public health infrastructure can lead to differential engagement in different countries. The U.S., in particular, warrants study because it leads the world in coronavirus infections and deaths (5) and the risk of adverse effects from COVID-19 vary substantially across groups, with Black and Hispanic Americans suffering disproportionately on these metrics (6).

Racial and ethnic disparities in COVID-19 health outcomes in the U.S. have frequently been attributed to feelings of mistrust toward the biomedical establishment rooted in a series of historic injustices (e.g., 7-9). Although mistrust may be a key factor shaping how members of minority groups engage with biomedical science during COVID-19 and other public health crises, the relationship is more complex. For instance, Black and Hispanic Americans have also long been underrepresented in biomedical science (e.g., 10-11), which limits science’s ability to serve members of these groups more generally—not just during a pandemic. This underrepresentation may reflect long-standing mistrust or even contribute to continued mistrust. Underrepresentation in turn can lead to negative experiences with medical care (12), institutionalized racism in health care (13-14), and systemic inequities in science that result from a majority white research community ill-positioned to engage with or pursue questions that are of central interest to other communities. Although mistrust is often a driving factor in the limited engagement in biomedical research among minority groups (e.g., 15-16), mistrust does not necessarily translate into lower willingness to participate in biomedical research (17-19), suggesting that additional factors may be at play. Moreover, defining the problem as one of communities mistrusting science risks ignoring the failures of the research community to effectively engage non-white communities.

This research seeks to shed light on the complexities that underlie how different subgroups within the general public, and particularly Black and Hispanic Americans, orient toward biomedical science. We begin with the observation that subgroups, including those who identify as a member of a racial and/or ethnic minority group, are too often treated as monoliths in behavioral and survey research—a tendency driven, in part, by the relatively small number of minority-identifying individuals that are included in a typical “representative” U.S. sample survey, relative to white individuals (e.g., 20). Although this broad brush approach may be helpful when trying to generate national-level estimates or between-group comparisons, we contend that it obscures a deeper understanding of how populations engage with the biomedical establishment. A more nuanced understanding of how different populations orient toward biomedical science is needed not only to inform strategies to engage different populations in health-promoting behaviors (e.g., vaccinations, mask-wearing), but also those aimed at encouraging participation in biomedical research, so that public health strategies are better able to serve diverse populations in the U.S. and around the world.

## Audience segmentation

Common approaches to understanding the public’s views toward science involve using demographic, political, or behavioral information (e.g., 21-23) or individual values dimensions (e.g., hierarchicalism/individualism; 24) to categorize people for analytic purposes. While such approaches frequently reveal valuable insights, people are not uni-dimensional; thus, approaches that categorize people on the basis of multiple dimensions may provide unique analytic insights. In this vein, drawing on techniques that originated in the social marketing literature, communication research suggests that audience segmentation analysis—an approach that categorizes individuals according to where they are placed on multiple relevant dimensions—can be fruitfully applied to understand how members of the public engage with a range of health-related issues (e.g., 25-26).

Applying an audience segmentation approach, we hypothesized that several dimensions related to values, motivations, and attitudes may simultaneously shape how Black, Hispanic, and general population U.S. adults engage with biomedical science. Because values-based segmentations cluster the population based on how similar they are on the hypothesized dimensions, this method may provide a clearer roadmap for scientists and other communicators to understand the public, connect to their values, and engage diverse audiences (e.g., 27-29).

As we detail below, through this process we identified seven dimensions that relate to a person’s orientation toward biomedical science based on previous literature as well as internal exploratory research. These dimensions include ones that capture attitudes toward science or biomedical science, as well as others that are not specifically science related but instead tap underlying values and worldviews. In other words, the segmentation conducted as part of this research incorporates various underlying values, beliefs, as well as attitudes toward biomedical science, but sets aside standard demographic and political categories in the segmentation. This approach allows for segmenting U.S. adult populations into smaller values-based groups, with the goal of offering more tailored insights for communication and engagement strategies.

This study pursued the following goals. First, in order to cluster each population on several dimensions simultaneously, we employed a novel application of a non-negative matrix factorization (NMF) model to a mindset segmentation. To accomplish this, we recruited not only a nationally representative sample of the U.S. adult general population, but two additional representative samples of Black and Hispanic adults in the U.S., to enable the research to better speak to the perspectives of communities that have been traditionally marginalized in biomedical research. Second, we sought to use these large and diverse samples as the basis for a segmentation analysis to increase understanding of the relationship between trust, concern, and biomedical science engagement between underserved populations and biomedical institutions. Finally, we sought to generate practical insights to help increase participation of underserved populations in biomedical research.

## Results

The segmentation analysis revealed a total of 10 distinct segments characterized by a unique attitudinal orientation toward biomedical science. More specifically, the analysis revealed five segments in each population (i.e., Black, Hispanic, and general population adults) with segment overlap, such that Segment 1 appears in all three populations, Segment 3 and Segment 5 appear in both the Hispanic and general populations, and Segment 4 appears in the Black and general populations. Figure 1 depicts each of the segments and their underlying dimensions.

**Figure 1.**
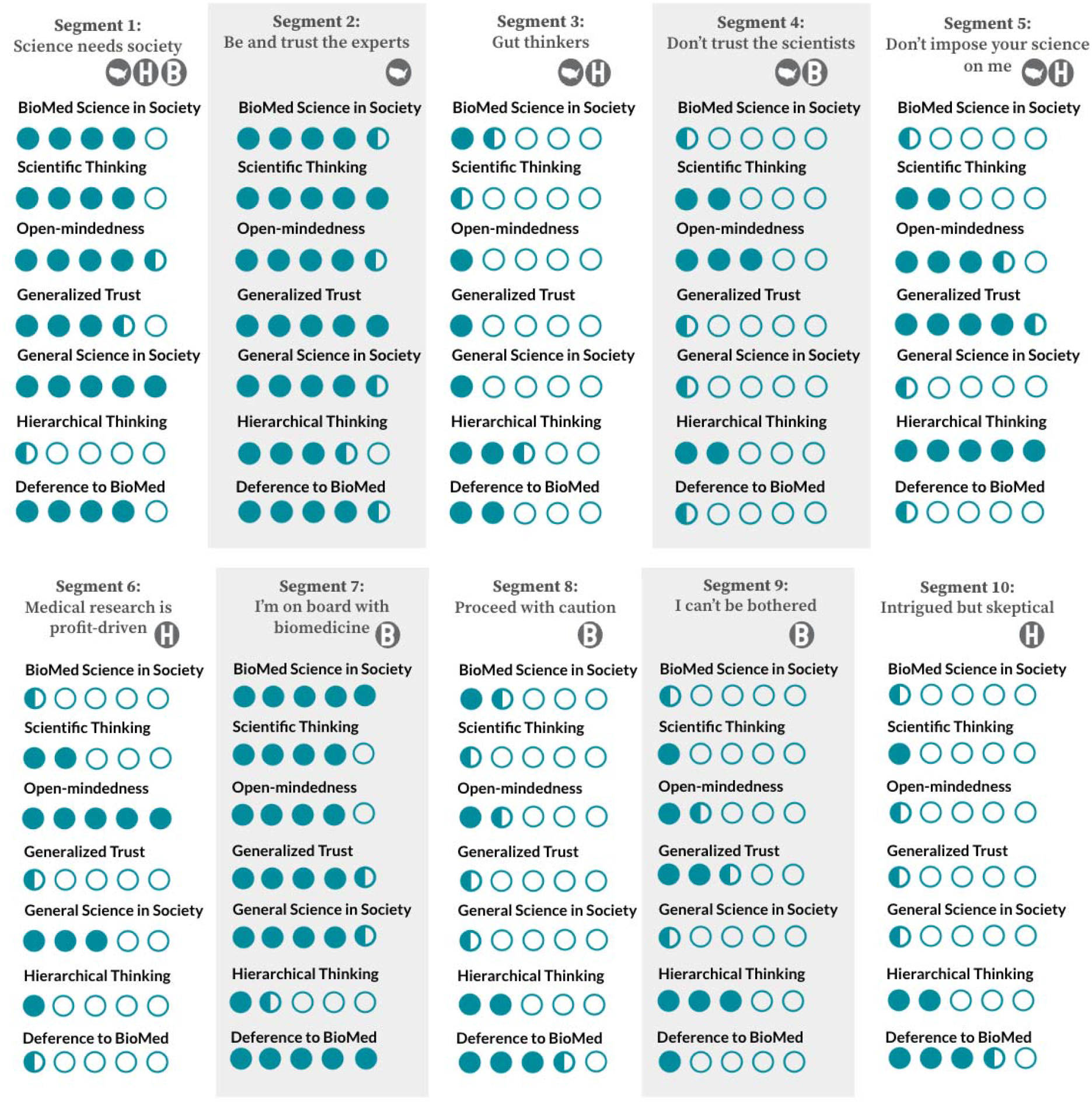
Overview of 10 biomedical mindset segments. Segments were identified based on how respondents answered 43 separate questions, each question associated with one of seven dimensions. The dots represent how well each dimension is reflected in each segment. Fewer dots indicate that the segment scored lower on a specific dimension, more dots indicate the segment scored higher on a specific dimension. An explanation for how the dots are calculated can be found in the Supplemental Material.

## The 10 biomedical science segments

Segment 1, which we’ve named *Society needs science*, is the only segment found in all three populations. It is the largest segment in the Hispanic (29%) and general populations (32%), and one of the smaller segments in the Black population (13%). This segment scores high on all dimensions except for hierarchical thinking, and especially high on Science in Society. The segment is characterized by a great deal of trust in science and the belief that both science and medical science should play a larger role in society, governmental decision-making, and engagement in their local community. Profiling variables that are represented to a greater degree in this population than average include that this group have doctors they can trust, believe that there is discrimination in biomedical institutions, and gathers information from left leaning or public news sources. Of all the segments, this has the highest proportion of college graduates and of people over 60.

Segment 2, *Be & trust the experts*, is found only in the general population sample (13%). This segment also scores high on all dimensions, and differs from Segment 1 for its relatively high reflection of hierarchical thinking. This segment not only trusts medical and scientific experts, but they also feel that they are the experts themselves and use scientific thinking in their everyday lives. They trust policymakers to use science and show high deference to biomedical authority. They are the youngest segment (containing the highest proportion of 18 to 29 year olds), are the least religious segment, and have a high proportion of college graduates.

Segment 3, labeled *Gut thinkers*, comprises a large proportion of both the general population (27%) and the Hispanic population (24%). This segment scores low on all dimensions, and especially low on scientific thinking. It is defined by a lack of interest in scientific thinking and a skepticism towards expertise, as well as a low desire to have science inform health policy. Gut thinkers generally lack curiosity and interest in scientific thinking, and do not believe that they possess scientific knowledge and may be more inclined to trust an internet search than a scientific authority, but turn to ABC or CBS for their news. This group is also disproportionately lower-income, lower-education, and generally does not have a medical professional that they can trust.

Segment 4, or *Don’t trust the scientists*, is found in both the general population sample (9%) and the Black sample (16%). They have low scores on most dimensions, but are relatively higher on open-minded thinking. The segment is characterized by a low generalized trust and low deference to biomedical authority and are least likely in the general population to have a doctor that they can trust. This segment is the most likely to have children in the home and has the highest proportion of people aged 30 to 44 years old. They tend to believe the seriousness of the COVID-19 pandemic has been exaggerated and prefer to get their news from “other” unspecified sources. In the Black population especially, this segment believes people in their community are treated poorly because of their race when seeking medical treatment.

Segment 5, *Don’t impose your science on me*, is found in both the general population sample (19%) and the Hispanic sample (19%). This group has high levels of generalized trust and holds individualistic views of society. They are generally skeptical of both governmental and scientific institutions and typically have low interest in science. They are mostly male, retired, and higher-income, and disproportionately turn to Fox News for their information. They are also the least likely to perceive any discrimination in biomedical institutions.

Segment 6, *Medical research is profit-driven*, is unique to the Hispanic sample (9%). This group believes science has a role to play in society and wants to increase science education in schools, yet believes that medical research is a tool of special interests. They are generally distrustful of others and have low deference to biomedical authority and are the segment in the Hispanic population with the lowest proportion of individuals who have gotten a regular checkup in the last year or to have a doctor they can trust. This segment has the highest proportion of people from Puerto Rico.

Segment 7, or *I’m on board with biomedicine*, is unique to the Black sample (28%). This group is defined by its high deference to biomedical authority and trusts them to make ethical decisions; they also tend to believe that science should inform health policy all of the time. This group has medical professionals they trust and have not experienced racial discrimination in the biomedical field. They are confident in their scientific knowledge and engage with science materials. They are curious and open minded, and prefer their news from left-leaning channels such as CNN and MSNBC. This group is mostly retired, male, older, educated, and higher-income.

Segment 8, *Proceed with caution*, is also unique to the Black sample (10%) and is generally distrustful of medical research and is low in general trust. Similar to segment 4, except for a relatively high score on the dimension for deference to biomedical authority. Even still, they remain concerned about people in their community being mistreated when seeking medical treatment because of their race, least likely among the segments in the Black population to have a doctor they can trust, and are also least likely to report having had a checkup in the last year. They turn to local news, ABC, or Fox for their information. This is a group that thinks the government interferes too much in society, and is mostly female, less educated, and has the highest proportion of lower-income earners of all segments.

Segment 9, *I can’t be bothered*, is also unique to the Black sample and is the largest segment in this population (34%). This group scores low on most dimensions, though is relatively higher in hierarchical thinking. The group is the most likely to have avoided seeking any medical treatment for fear of racial discrimination, and they are unlikely to have a medical professional that they can trust. They place little trust in science and show low deference to biomedical authority. They also are low engagement with regards to scientific thinking, indicating some detachment from science. This group is also disproportionately unemployed.

Segment 10, or *Intrigued but skeptical*, is unique to the Hispanic sample (19%). This group clustered together in part because they appear to exhibit high acquiescence bias. Even though removing straightliners was part of our quality control measures, this group was extremely agreeable to most segmentation questions including contradictory statements, which is a known risk with agree/disagree statements (e.g., 30). Moreover, this segment is disproportionately lower-income and a plurality (45%) did not graduate from high school, which are characteristics consistent with satisficing and/or acquiescence. Although this group did emerge as a segment, given this potential for bias, we cautiously elected not to include them in the analysis reported below, pending additional qualitative research to more fully understand this group.

## Relationship between trust and segment

As expected, trust emerged as a significant predictor of segment assignment; thus, we elected not to include it as a covariate in our regression models (see Table 1).

**Table 1:** Segment Assignment Predicted by Trust, Multinomial Logistic Regression.

|  |  | Model 1.<br>General Population |  | Model 2.<br>Hispanic Population |  | Model 3.<br>Black Population |  |
| --- | --- | --- | --- | --- | --- | --- | --- |
| Segment |  | Intercept | Trust Count | Intercept | Trust Count | Intercept | Trust Count |
| 2 | Coef | -0.69 (0.37) | -0.05 (0.09) |  |  |  |  |
|  | Risk Ratio | 0.50 | 0.95 |  |  |  |  |
| 3 | Coef | 1.85<sup>***</sup> (0.26) | -0.58<sup>***</sup> (0.07) | 1.84<sup>***</sup> (0.26) | -0.59<sup>***</sup> (0.07) |  |  |
|  | Risk Ratio | 6.35<sup>***</sup> | 0.56<sup>***</sup> | 6.28<sup>***</sup> | 0.55<sup>***</sup> |  |  |
| 4 | Coef | 1.22<sup>***</sup> (0.30) | -0.76<sup>***</sup> (0.09) |  |  | 1.13<sup>***</sup> (0.26) | -0.29<sup>***</sup> (0.07) |
|  | Risk Ratio | 3.39<sup>***</sup> | 0.47<sup>***</sup> |  |  | 3.10<sup>***</sup> | 0.75<sup>***</sup> |
| 5 | Coef | 2.06<sup>***</sup> (0.26) | -0.79<sup>***</sup> (0.07) | 2.13<sup>***</sup> (0.26) | -0.80<sup>***</sup> (0.07) |  |  |
|  | Risk Ratio | 7.83<sup>***</sup> | 0.45<sup>***</sup> | 8.42<sup>***</sup> | 0.45<sup>***</sup> |  |  |
| 6 | Coef |  |  | 1.29<sup>***</sup> (0.29) | -0.77<sup>***</sup> (0.09) |  |  |
|  | Risk Ratio |  |  | 3.65<sup>***</sup> | 0.46<sup>***</sup> |  |  |
| 7 | Coef |  |  |  |  | -1.74<sup>***</sup> (0.39) | 0.63<sup>***</sup> (0.09) |
|  | Risk Ratio |  |  |  |  | 0.17<sup>***</sup> | 1.88<sup>***</sup> |
| 8 | Coef |  |  |  |  | -0.45 (0.35) | 0.06 (0.09) |
|  | Risk Ratio |  |  |  |  | 0.64 | 1.06 |
| 9 | Coef |  |  |  |  | 1.74<sup>***</sup> (0.25) | -0.24<sup>***</sup> (0.07) |
|  | Risk Ratio |  |  |  |  | 5.68<sup>***</sup> | 0.79<sup>***</sup> |
|  | Residual Deviance | 2698.05 |  | 1850.62 |  | 2690.82 |  |
|  | AIC | 2714.047 |  | 1862.62 |  | 2706.82 |  |
|  | N | 957 |  | 780 |  | 963 |  |

\* $p < .05$ , \*\* $p < .01$ , \*\*\* $p < .001$
Note: Standard errors in parentheses
The table includes estimates of unstandardized coefficients from three separate multinomial logistic regression models predicting segment assignment from trust count, one for each of the three adult populations of interest: general population, Black population, Hispanic population. Risk ratios are presented for each model. Similar in interpretation to odds ratios, the risk ratio refers to the odds of being more likely to belong to that segment, compared to the reference category (Segment 1).

## Relationship between concerns and engagement

Overall, results reveal that willingness to engage is above 10% for each population for each type of volunteer activity (see Figure 2). On average, we find that 17% (SE = 1.06%) of adults are extremely or very likely to be willing to participate in medical research for the Centers for Disease Control if given the opportunity; 12% (SE = 0.93%) for a vaccine trial at a local university; 17% (SE = 1.08%) for donating tissue to a local university for medical research; 16% (SE = 1.06%) for participating in medical research at the CDC; 15% (SE = 0.96%) for donating money to support medical research. There were no significant differences among the populations with respect to willingness to volunteer. In other words, an average adult from the General Population sample was just as likely to be willing to volunteer for each activity as an average adult from the Black or Hispanic samples.

**Figure 2:**
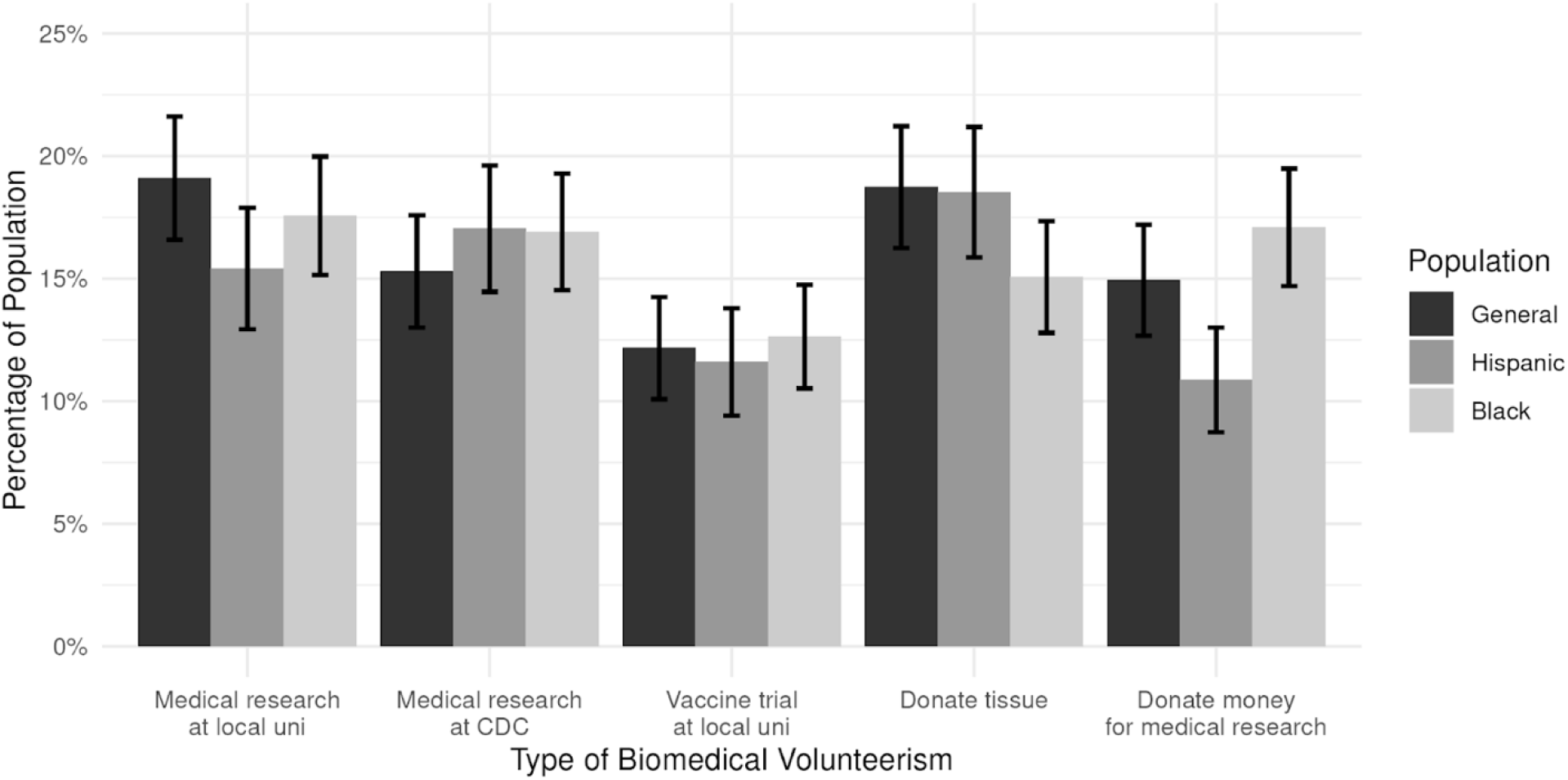
Average willingness to participate in medical research by population. The height of the bar indicates the percentage of adults in each population that is very or extremely likely to do each of the activities if given the opportunity.

In addition, and in line with previous research (e.g., 31), we find higher levels of concerns about participating in biomedical research among the Black and Hispanic samples as compared to the general population sample, which held for all concerns except for having to pay. Compared to the general population sample, the Black and Hispanic samples are significantly more concerned about being a guinea pig (Black adults: d=0.14, p<0.001; Hispanic adults: d=0.07, p<0.05), getting COVID-19 (Black adults: d=0.21, p<0.001; Hispanic adults: d=0.12, p<0.001), getting a disease other than COVID-19 (Black adults: d=0.15, p<0.001; Hispanic adults: d=0.15, p<0.001), privacy (Black adults: d=0.14, p<0.001; Hispanic adults: d=0.11, p<0.01), trustworthiness of the researcher and the process (Black adults: d=0.14, p<0.001; Hispanic adults: d=0.12, p<0.001) (see Figure 3).

**Figure 3.**
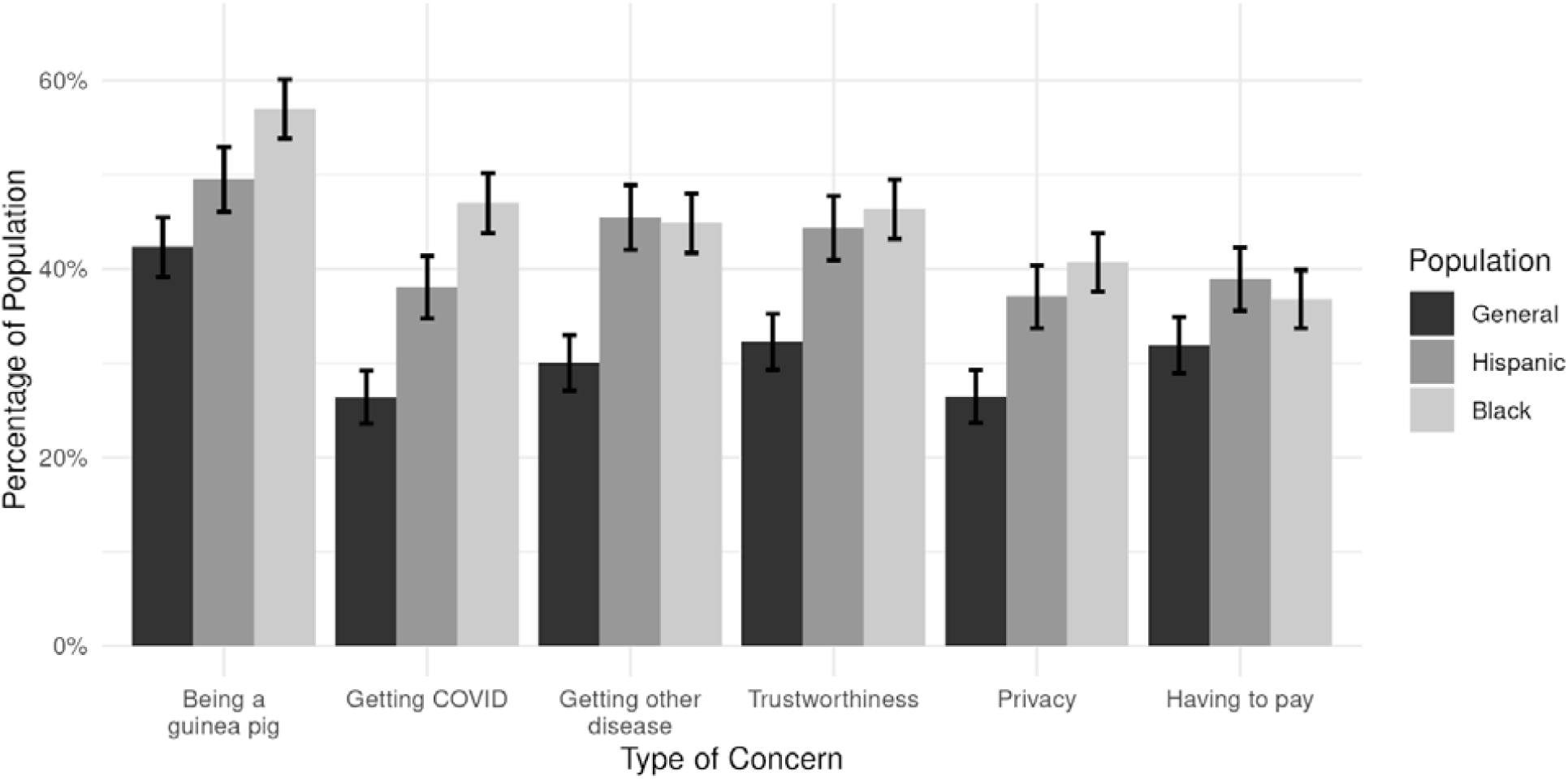
Average concern by population. The height of the bar indicates how much each of the concerns is “extremely” or “very” concerning to each population, on average. Black and Hispanic adult populations consistently have higher concerns.

However, we also find that the relationship between concerns and willingness to participate in biomedical research varied across the three samples. Whereas concerns were a significant negative predictor of willingness to participate for four of the five activities we asked about among the general population (Bs: -.10 to -.22, ps < .01) and Hispanic samples (Bs: -.10 to -.17, ps < .01), this relationship did not emerge for Black adults. In other words, the more concerns that a general population or Hispanic adult expressed, the less willing they were to participate, which held for every volunteer activity we asked about except donating money. By comparison, the only negative relationship that emerged between concerns and willingness to participate within the Black sample was for participating in a vaccine trial at a university (B: -.08, p < .05) (see Tables 2-4 for complete list of regression coefficients and odds ratios for the general population, Hispanic population, and Black sample, respectively).

**Table 2:** Biomedical Science Volunteerism by Segment and Concern, General Population Ordinal Logistic Regression.

| Variable |  | Participate in Medical Research at University | Participate in Medical Research for CDC | Participate in Vaccine Trial at University | Donate Tissue at University | Donate Money to Medical Research |
| --- | --- | --- | --- | --- | --- | --- |
| Concern Count | *Coef* | -0.14*** (0.04) | -0.11** (0.04) | -0.22*** (0.04) | -0.10** (0.04) | -0.00 (0.03) |
|  | *OR* | 0.86*** | 0.89** | 0.80*** | 0.90** | 0.99 |
| Segment 2 | *Coef* | 0.35 (0.20) | 0.03 (0.20) | -0.04 (0.21) | 0.21 (0.20) | 0.00 (0.20) |
|  | *OR* | 1.42 | 1.03 | 0.96 | 1.23 | 1.00 |
| Segment 3 | *Coef* | -0.82*** (0.17) | -0.88*** (0.17) | -0.86*** (0.19) | -0.69*** (0.17) | -0.86*** (0.17) |
|  | *OR* | 0.44*** | 0.41*** | 0.42*** | 0.50*** | 0.42*** |
| Segment 4 | *Coef* | -0.24 (0.24) | -0.22 (0.23) | -0.33 (0.26) | -0.57* (0.24) | -0.60* (0.24) |
|  | *OR* | 0.79 | 0.80 | 0.72 | 0.56* | 0.55* |
| Segment 5 | *Coef* | -1.34*** (0.20) | -1.78*** (0.22) | -1.98*** (0.28) | -1.04*** (0.19) | -1.32*** (0.19) |
|  | *OR* | 0.26*** | 0.17*** | 0.14*** | 0.35 | 0.27*** |
| *AIC* |  | 2026.07 | 1946.137 | 1616.65 | 2082.34 | 2020.34 |
| *N* |  | 957 | 957 | 957 | 957 | 957 |

\* $p < .05$ , \*\* $p < .01$ , \*\*\* $p < .001$
Note: Standard errors in parentheses
The table includes estimates of unstandardized coefficients from five ordinal logistic regression models predicting willingness to participate in a specific type of medical research (columns), from concern\_count, controlling for segment for the general population. Odds ratios (OR) are presented for each model. The reference category is Segment 1.

**Table 3:** Biomedical Science Volunteerism by Segment and Concern, Hispanic Population Ordinal Logistic Regression.

| Variable |  | Participate in Medical Research at University | Participate in Medical Research for CDC | Participate in Vaccine Trial at University | Donate Tissue at University | Donate Money to Medical Research |
| --- | --- | --- | --- | --- | --- | --- |
| Concern Count | *Coef* | -0.13*** (0.04) | -0.10** (0.04) | -0.17*** (0.04) | -0.12*** (0.04) | -0.07 (0.04) |
|  | *OR* | 0.88*** | 0.90** | 0.84*** | 0.89*** | 0.93 |
| Segment 3 | *Coef* | -1.39*** (0.19) | -1.00*** (0.19) | -1.13*** (0.21) | -0.77*** (0.18) | -1.22*** (0.19) |
|  | *OR* | 0.25*** | 0.37*** | 0.32*** | 0.46*** | 0.30*** |
| Segment 5 | *Coef* | -1.53*** (0.20) | -1.46*** (0.20) | -1.48*** (0.23) | -1.09*** (0.19) | -1.59*** (0.20) |
|  | *OR* | 0.22*** | 0.23*** | 0.23*** | 0.33*** | 0.20*** |
| Segment 6 | *Coef* | -0.03 (0.23) | 0.07 (0.24) | -0.73** (0.27) | -0.10 (0.23) | -0.72** (0.24) |
|  | *OR* | 0.97 | 1.08 | 0.48** | 0.90 | 0.49** |
| *AIC* |  | 1585.72 | 1600.78 | 1318.82 | 1676.81 | 1490.08 |
| *N* |  | 780 | 780 | 780 | 780 | 780 |

\* $p < .05$ , \*\* $p < .01$ , \*\*\* $p < .001$
Note: Standard errors in parentheses
The table includes estimates of unstandardized coefficients from five ordinal logistic regression models predicting willingness to participate in a specific type of medical research (columns), from concern\_count, controlling for segment for the Hispanic population. Odds ratios (OR) are presented for each model. The reference category is Segment 1.

**Table 4:** Biomedical Science Volunteerism by Segment and Concern, Black Population Ordinal Logistic Regression.

| Variable |  | Participate in Medical Research at University | Participate in Medical Research for CDC | Participate in Vaccine Trial at University | Donate Tissue at University | Donate Money to Medical Research |
| --- | --- | --- | --- | --- | --- | --- |
| Concern Count | *Coef* | -0.01 (0.03) | -0.00(0.03) | -0.08* (0.03) | -0.02(0.03) | 0.09** (0.03) |
|  | *OR* | 0.99 | 1.00 | 0.92* | 0.98 | 1.09** |
| Segment 4 | *Coef* | -1.12*** (0.23) | -0.92*** (0.24) | -0.73** (0.27) | -0.93*** (0.24) | -0.36 (0.23) |
|  | *OR* | 0.33*** | 0.40*** | 0.48** | 0.39*** | 0.69 |
| Segment 7 | *Coef* | -0.43* (0.20) | -0.09 (0.20) | -0.08 (0.22) | -0.60** (0.21) | -0.05 (0.20) |
|  | *OR* | 0.65* | 0.92 | 0.92 | 0.55** | 0.94 |
| Segment 8 | *Coef* | -1.21*** (0.28) | -1.01*** (0.28) | -0.36 (0.30) | -0.74** (0.27) | -0.72** (0.26) |
|  | *OR* | 0.30*** | 0.36*** | 0.70 | 0.48** | 0.48** |
| Segment 9 | *Coef* | -0.67*** (0.20) | -0.54** (0.20) | 0.16 (0.22) | -0.65** (0.20) | -0.25 (0.20) |
|  | *OR* | 0.51*** | 0.58*** | 1.17 | 0.52*** | 0.78 |
| *AIC* |  | 2141.74 | 2083.94 | 1774.80 | 2003.67 | 2165.95 |
| *N* |  | 963 | 963 | 963 | 963 | 963 |

\* $p < .05$ , \*\* $p < .01$ , \*\*\* $p < .001$
Note: Standard errors in parentheses
The table includes estimates of unstandardized coefficients from five ordinal logistic regression models predicting willingness to participate in a specific type of medical research (columns), from concern\_count, controlling for segment for the Black population. Odds ratios (OR) are presented for each model. The reference category is segment 1.

## “Being a guinea pig” as a leading concern

We were also interested in which concerns would emerge as the strongest predictor of engagement across the samples. Taken together, fear about being a guinea pig was the most consistent and negative predictor of engagement across all three populations (Table 5). Furthermore, this concern remained a significant and negative predictor when controlling for the various segments that emerged across all three populations (see Table 6). Finally, our regression analysis found no evidence of a significant interaction between fear of being a guinea pig and segment when predicting engagement.

**Table 5:** Effect of Concerns on Willingness to Participate in Biomedical Research by <u>Population, Ordinary Least Squares Regression</u>.

| Concern | Model 1. | Model 2. | Model 3. |
| --- | --- | --- | --- |
|  | General Population | Hispanic Population | Black Population |
| Being a “Guinea Pig” | -2.40*** (0.26) | -2.28*** (0.28) | -1.45*** (0.29) |
| Trustworthiness of the Researcher | -0.22 (0.27) | -0.14 (0.28) | -0.33 (0.28) |
| Getting Covid | 0.37 (0.29) | 0.10 (0.29) | 0.45 (0.28) |
| Getting a Disease Other than Covid | 0.56 (0.30) | -0.59* (0.30) | 0.47 (0.31) |
| Results Not Confidential | -0.42 (0.28) | 0.19 (0.27) | -0.27 (0.28) |
| Having to Pay for Treatments | 0.80** (0.26) | 1.04*** (0.26) | 1.39*** (0.27) |
| Intercept | 8.86*** (0.16) | 9.08*** (0.18) | 8.43*** (0.19) |
| *F-Statistic* | 18.71 | 21.01 | 10.22 |
| *Adj. R<sup>2</sup>* | 0.10 | 0.13 | 0.06 |
| *N* | 957 | 780 | 963 |

\* $p < .05$ , \*\* $p < .01$ , \*\*\* $p < .001$
Note: Standard errors in parentheses

**Table 6:** Effect of “Guinea Pig” Concern on Willingness to Participate in Biomedical Research by Population, Ordinary Least Squares Regression.

|  | Model 1.<br>General<br>Population | Model 2.<br>Hispanic<br>Population | Model 3.<br>Black<br>Population |
| --- | --- | --- | --- |
| “Guinea Pig”<br>Concern | -1.84<sup>***</sup> (0.22) | -1.81<sup>***</sup> (0.22) | -0.72<sup>**</sup> (0.23) |
| Segment 2 | 0.65 (0.35) |  |  |
| Segment 3 | -1.37<sup>***</sup> (0.28) | -1.84<sup>***</sup> (0.28) |  |
| Segment 4 | -0.34 (0.40) |  | -1.60<sup>***</sup> (0.43) |
| Segment 5 | -2.16<sup>***</sup> (0.31) | -2.33<sup>***</sup> (0.29) |  |
| Segment 6 |  | -0.23 (0.38) |  |
| Segment 7 |  |  | -0.60 (0.38) |
| Segment 8 |  |  | -1.54<sup>**</sup> (0.48) |
| Segment 9 |  |  | -0.95<sup>*</sup> (0.37) |
| Intercept | 9.69<sup>***</sup> (0.19) | 10.15<sup>***</sup> (0.19) | 9.58<sup>***</sup> (0.34) |
| *Adj. R*<sup>2</sup> | 0.16 | 0.20 | 0.03 |
| *N* | 957 | 780 | 963 |

\* $p < .05$ , \*\* $p < .01$ , \*\*\* $p < .001$
Note: Standard errors in parentheses
The table provides estimates for unstandardized coefficients from three different ordinary least squares regression models, one for each population (column). The predictor variable is guinea pig concern which are coded as 1 (“very” or “extremely” concerned) or 0 (reference category). The dependent variable is an engagement score which ranges from 5-20 and is created by combining each of the five engagement variable (donate tissue, medical research at the CDC, etc.) scores which takes on a value of 1 (extremely unlikely) to 4 (extremely likely). Being a guinea pig is a consistent negative predictor across all populations (including when controlling for segment).

## Discussion

Better understanding how different groups engage with biomedical science remains a critical priority for outreach efforts aimed at improving public health outcomes, particularly among groups that have been historically mistreated and marginalized by the biomedical establishment. While some have suggested that the underrepresentation of racial and ethnic minority communities in medical research is due to their lack of willingness to participate (e.g., 32), other work challenges this notion, by demonstrating that minorities’ greater concerns or fears about participation do not necessarily translate in less willingness to engage (17-18) and that consent rates do not always differ by race (19). The present research extends these findings by taking a closer look at the factors that shape engagement with biomedical research among three independent and nationally representative samples of U.S. adults–specifically, a general population sample, a sample of Hispanic adults, and a sample of Black adults. By analyzing dedicated samples of these three distinct populations, our study is able to provide a deeper level of nuance than most single-sample survey studies, which often do not include sufficient samples of Black and Hispanic adults to provide for meaningful analysis, which limits the insights that can be drawn about key populations that public health efforts are keen to understand.

Our unique datasets and mindset segmentation approach, which categorizes members of the public based on multiple values-based dimensions simultaneously, revealed a number of important findings. Importantly, we replicate earlier findings (17-19) that despite expressing greater concerns about participating in biomedical research, this does not result in differential willingness to participate across Black, Hispanic, or general population adults in the U.S. Interestingly, however, we also find that while concerns was a negative predictor of engagement in biomedical research among general population and Hispanic adults, this was not the case among Black adults, which counters the prevailing notion that longstanding fears rooted in mistrust toward the biomedical establishment are causing this group to disengage. Moreover, our analysis of leading concerns found that one concern stood out as a strong and negative predictor of engagement for all groups: the fear of being a guinea pig, which predicted lower willingness to volunteer in each activity we tested, except for donating money, which held not only across the three survey samples but across each of the 10 segments that emerged in our analysis. Given the consistency of this fear as a negative predictor across all populations and segments, future research may wish to explore what being a guinea pig means to the different segments, as well as potential ways to reduce this fear, especially among groups that are disproportionately affected by illness and disease and that would stand the benefit the most from greater engagement with medical science.

This study also builds on previous research by furthering our understanding of the nuance in how these different populations orient towards biomedical science. The values-based mindset segmentation we employed here allowed us to identify sub-segments of each population based on various beliefs and values regarding how the world works, how the world should work, and the role that science and biomedical science should play in society. By revealing both universal segments that emerged across populations and segments that were unique to particular populations, our segmentation analysis reveals underappreciated heterogeneity across our samples. Stated differently, Black and Hispanic adults are not monolithic, and by more closely examining the variation in how individuals within these groups orient toward biomedical science, medical researchers and others working on community engagement are better positioned to design effective strategies for engaging the public.

Of the 10 mindsets identified in the U.S. population, one is unique to the general population, two are unique to the Hispanic adult population, and three are unique to the Black adult population. The unique segments in the general population (Segment 2: Be & trust the experts) and Black populations (Segment 7: I’m on board with biomedicine) include individuals that are highly deferential to biomedical science. Additional unique segments in the Black population include a skeptical segment (Segment 8: Proceed with caution) and a detached segment (Segment 9: I can’t be bothered). The unique segments in the Hispanic adult population include one that is supportive and interested in science generally, but is somewhat skeptical and cautious about medical science specifically (Segment 6: Medical Research is profit-driven). The skeptical segments do not indicate less willingness to engage, but rather appear to require trust-building in order to feel comfortable engaging.

When viewed through the lens of values-based segments rather than through the more familiar lens of demographic variables (e.g., Black, Hispanic), a more nuanced picture emerges, with each population containing segments that are interested and willing to engage with biomedical science if given the opportunity, while also containing segments that distrust science or biomedical institutions and that are detached from science generally. Segments 1 (Society needs science), 2 (Be & trust the experts), and 7 (I’m on board with biomedicine) are the most trusting and interested in biomedical science, and together these segments make up 43% of the general population, 29% of the Hispanic population, and 41% of the Black population. Yet, if we look at data on medical research participation, it is overwhelmingly represented by people of European ancestry. For example, Gurdsani and colleagues (33) find that the proportion of individuals in genome studies that are critical to understanding genetic determinants of disease are overwhelmingly of European heritage (78%). In addition, Nazha and colleagues (34) caution that the disproportionately low proportion of minority patients in immunotherapy clinical trials could lead to poorer outcomes for those communities. As Katz and colleagues (17-18) suggest, this underrepresentation may reflect that individuals from underrepresented communities have not had the opportunity to participate, rather than their level of willingness to engage.

As with any mindset-based segmentation, there is an element of researcher choice in deciding on the optimal segmentation solution. The process involves considering various model metrics and thinking about segments with practical fidelity in mind. Another research team may conclude a different segmentation outcome. Thus, this research would benefit from additional research further validating the findings and interpretation of the results. Qualitative research including interviews or focus groups with representatives of these various segments would increase the understanding of how to engage each population, given the various concerns, barriers and lived experiences unique to each mindset.

While the timing of these surveys (October-November 2021) had the benefit of allowing us to analyze these different mindsets during a historically significant biomedical event (the COVID-19 pandemic), the broader societal conversations regarding trust in science and social disparities in medical outcomes occurring at that time may well have influenced the patterns we observed here. Future research should attempt to replicate these findings, to examine whether they remain with greater temporal distance from the start of the pandemic.

Understanding how different groups in society view biomedical science is widely recognized as a key factor in addressing persistent public health disparities; yet, efforts to further this understanding too often rely on survey samples that are too small to enable subgroup analysis, or otherwise employ methods that are not well suited to exploring the heterogeneity of views that exist within groups that have been traditionally marginalized in biomedical science, such as Black and Hispanic individuals in the United States. We addressed these limitations by recruiting independent nationally representative samples of Black, Hispanic, and general population U.S. adults and by employing a non-negative matrix factorization model in a novel way to conduct a values-based mindset segmentation analysis. This revealed significant variation both within and between these populations with regard to their views toward the biomedical establishment.

Our findings may inform efforts aimed at increasing representation in medical research, which is one step in addressing persistent racial and ethnic health disparities. Within some populations, there may be a long way to go, given that a majority remains science-skeptical or science-detached (57% of the general population, 71% of the Hispanic population, 59% of the Black population). These segments are less trusting of biomedical institutions and professionals, and therefore may be less willing to accept the findings of medical research. However, there remains a significant opportunity to deepen relationships with sizable segments interested and willing to engage with biomedical research (45% of the general population, 29% of the Hispanic population, and 41% of the Black population). Sustaining engagement with these segments will likely require intentional efforts of relationship building between the biomedical field and under-represented populations in the short-term, and more systemic changes (e.g., increasing representation in biomedical science, sustaining support for researchers and communities to co-develop knowledge and interventions) over the long-term. These efforts can encourage trust in the biomedical field, such that various populations will engage with research findings and recommendations drawn from more representative data. Future work should incorporate a nuanced understanding of the beliefs, concerns, and values across communities, in order to inform a suite of effective approaches to meet people where they are at and build a more trustworthy scientific enterprise.

## Materials and Methods Sample

We administered three national probability sample surveys, one for each of the following adult populations: Black adults in the U.S. (N= 963), Hispanic adults in the U.S. (N=964), and the general population of U.S. adults (N=957). The surveys were fielded from October 21, 2021, to November 6, 2021, on the AmeriSpeak panel maintained by NORC at the University of Chicago, which uses probability-based methods to recruit representative samples of respondents. Respondents classified as rushers (those who completed the survey in less than ⅓ of the median completion time), straightliners (respondents who straightlined responses on three or more segmentation battery grids), or skippers (respondents who skipped more than 50% of total questions) were removed from the sample (N = 126). NORC applied a panel base sampling weight to account for probability of selection into the panel, to align with U.S. Census benchmarks for each, and to account for potential nonresponse bias among AmeriSpeak panelists. Post-stratification weights were applied to account for any deviations from the Census benchmarks for each population.

## Measures

### Segmentation battery

We designed a 43-item battery of questions to assess each of the seven dimensions that we hypothesized *a priori* to underlie the public’s engagement with biomedical science. Survey questions were either adapted from previous research (i.e., the items for *Curiosity and open-minded thinking, Hierarchical/egalitarian, Generalized trust*, and *Deference to scientific authority*; see citations below) or were created based on internal exploratory research (i.e., the items for *Scientific thinking, Science in society*, and *Biomedical science in society*). While the dimensions are a useful means of organizing the segmentation battery and in synthesizing the NMF model output, these *a priori* dimensions do not factor into the NMF clustering model input. The complete list of battery questions can be found in the Supplemental Material.

#### Curiosity and open-minded thinking (8 items)

This dimension was intended to capture curiosity and open-minded thinking, which research suggests can counteract polarization in the interpretation of scientific information (e.g., 35). This dimension provides an indication of how individuals update their understanding based on new information as well as an indication of one’s susceptibility to confirmation bias. Example survey items include, “I consult multiple sources to make difficult decisions” and “I prefer news that aligns with or supports my beliefs” (reverse-scored).

#### Hierarchical thinking (9 items)

This dimension was intended to capture one’s view of how the social world should function and how society should be organized, which has been found to predict how audiences assess risk and act on scientific information (e.g., 36). Example survey items include, “The government interferes far too much in our everyday lives” (reverse-scored) and “Respect for authority is something all children need to learn.”

#### Generalized trust (7 items)

This dimension was intended to capture one’s level of generalized trust in others and society, which can influence a person’s engagement in civic activities and possibly one’s willingness to engage with research for the public good (37-38). Example survey items include, “Generally speaking, I trust other people” and “Generally speaking, other people treat me unfairly” (reverse-scored).

#### Scientific thinking (5 items)

This dimension was intended to capture one’s use and understanding of science in their everyday lives, which is expected to predict willingness to engage with biomedical research and science in general (39). Example survey items include, “The science education I received in school helps me in my everyday life” and “I trust my gut more than experts when making important decisions” (reverse-scored).

#### Science in society (8 items)

This dimension was intended to capture the role people perceive science should play in society and policy-making. The more a person believes science should play a role in society, the more willing they may be to participate in science. However, people may perceive biomedical research and scientific research separately and have different views about the role each should play in society (we therefore included a dimension specifically on biomedical science in society as the next dimension). Example survey items include, “I’d pay more taxes to fund scientific research that directly benefits my local community” and “Science is just another tool of manipulation that those in power can use to push their agenda” (reverse-scored).

#### Role of biomedical science in society (6 items)

This dimension was intended to capture the role people perceive biomedical science, specifically, should play in society and policy-making. Americans who believe biomedical science fundamentally improves lives and that it should inform public policy are expected to be more willing to engage in biomedical science. Example survey items include, “Medical science creates just as many problems as it solves” (reverse-scored) and “The work of medical science researchers helps my community.”

#### Deference to biomedical authority (5 items)

This dimension was adapted from previous research (40) and was intended to capture deference to biomedical science authority, which is expected to relate to Americans’ beliefs about how science-related decisions should be made and their personal engagement with biomedical research. Example items include, “Medical doctors have our best interest in mind when they tell the public what to do” and “I trust medical researchers to make ethical decisions.”

### Main outcome variables

The following three variables comprised the main outcome variables of our segmentation analysis:

#### Trust in biomedical professionals and institutions

Respondents were asked, “How much do you trust each of the following to do what is right for you or your family?” and were provided the following five entities in random order: *University medical research center, Primary health care provider (e*.*g*., *local health care clinic, primary care office), Local public health agency, The federal government (Centers for Disease Control, National Institute of Health, Federal Drug Administration)*, and *A pharmaceutical company*. The response options were “A great deal,” “A fair amount,” “Not too much,” and “No trust at all.” The resulting variable, “trust_count,” ranged from 0-5, and incremented each time a respondent answered “A great deal” or “A fair amount.”

#### Engagement in medical research opportunities

We define engagement here as willingness to volunteer in medical research. Respondents were asked, “How likely are you to do any of the following activities, if the opportunity arises?” and were provided the following five opportunities in random order: *Participate in medical research at a local university, Participate in a vaccine trial at a local university, Donate tissue to a local university for medical research, Participate in medical research for the federal Centers for Disease Control*, and *Donate money to support medical research*. The response options were “Extremely likely,” “Very likely,”, “Somewhat likely,” and “Not at all likely,” and were coded as ordinal (4, 3, 2, and 1, respectively) to enable modeling the degree of engagement.

#### Concerns about engaging in volunteer activities

In this battery of questions adopted from Katz and colleagues (17), respondents were asked, “There are lots of reasons some people do **NOT** want to participate in medical research studies. When you think about participating in medical research studies, how concerned are you about each of the following?” and were provided the following six concerns in random order: *Getting COVID-19, Getting a disease other than COVID-19, Being a ‘guinea pig*,*’ Results not being private or confidential, Having to pay for research treatments*, and *The trustworthiness of the research process or the researcher*. The response options were “Extremely concerned,” “Very concerned,” “Moderately concerned,” “Slightly concerned,” and “Not at all concerned.” The resulting variable, “concern_count,” ranged from 0-6, and incremented each time a respondent answered “Very” or “Extremely concerned.”

#### Additional variables that describe the segments

In addition to the segmentation battery, the survey included additional questions to further characterize the different segments. These variables appear in the Supplemental Material (section 3) and include questions related to experiences with biomedical institutions, engagement in the community, sources of news, and engagement with various scientific information.

## Analytic approach

### Segmentation analysis

Our segmentation analysis sought to address the following questions: Into how many groups can each population be divided, and what are their sizes? What are the unique and defining characteristics that describe each of the mindset segments? Prior research suggests that non-negative matrix factorization (NMF) is best equipped to address these questions (41-43). NMF can be used for several applications of multivariate data and has a benefit over other clustering algorithms in that it is not subject to the curse of dimensionality like other algorithms such as K-Means or Hierarchical clustering (44-45). Additionally, NMF has become a widely used tool for the analysis of high dimensional data as it automatically extracts sparse and meaningful features from a set of nonnegative data vectors. NMF’s use in segmentation analysis is rare; it is more typical for segmentations to utilize either demographic characteristics or be constrained by a different clustering model that requires fewer dimensions (and thus fewer segmentation battery questions).

The NMF algorithm produces probabilities (e.g., the probability of being assigned to a certain segment), and the respondent is assigned to the segment with the highest associated probability. The final segmentation determination was made by evaluating the performance of the model by checking the dispersion, silhouette and sparseness scores, and a generic size breakdown for the given K’s. It is also important to note that there is not necessarily only one solution, and that multiple characteristics of each solution are evaluated against each other to select the best fit. For more details on this approach, see the Supplemental Material.

Once the populations are clustered into segments, it is necessary to understand how each dimension is reflected in each segment. To do this, question-specific index values were used. An index value is the percentage of respondents in a segment that agreed (“strongly agree” or “somewhat agree”) with the statement divided by the overall agreement of all respondents times 100 (see Equation 1). An index value of 100 indicates no differentiation between the segment and the overall population, while index values above 100 and below 100 indicate that the segment agrees more or less with the statement compared to the general population, respectively. The distance from 100 was then calculated, and the distance was multiplied by -1 to account for questions that are reverse coded.

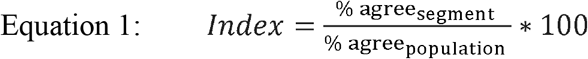

Because each question was part of a series of questions in a dimension battery, the mean distances for all questions in a dimension were calculated for each segment. All means were then scaled from 0-1, with 0 corresponding to the lowest value for each category and 1 corresponding to the highest value in each category. This enables us to understand how the different dimensions comprise the segments, but also helps to differentiate between the segments, and were the basis for the dots in Figure 1.

### Regression methods

To uncover additional insights that can inform engagement strategies, we explored the relationship between and among segments, trust, engagement and concern through the following analyses.

#### Trust and segment

Given that the segments are based on various underlying dimensions that include trust in biomedical institutions (i.e., generalized trust and deference to biomedical authority), we expect that trust in different medical institutions will be highly predictive of segment assignment. If this is the case, we would not want to include both trust and segment as covariates in a regression model predicting engagement. To evaluate this hypothesis, we estimated parameters from three multinomial logistic regressions (one for each population) predicting segment assignment from *trust_count* with Segment 1 (*Society needs science*; see full list below) as the reference category

#### Engagement

To understand if concerns impact willingness to participate in different medical activities, we first estimated parameters of 15 ordinal logistic regressions predicting each volunteer activity (five total) from segment plus *concern_count* for each population (three total). The dependent variable is ordinal; it is based on a scale ranging from “Not at all likely” to “Extremely likely” to participate. Thus, an ordinal logistic regression model is appropriate to assess each outcome. The segment was a dummy variable and Segment 1 – which was present in all three surveyed populations – was the omitted category.

#### Leading concerns

Finally, to understand whether there were any leading concerns that predicted engagement across all populations, we estimated parameters of three OLS regressions predicting total engagement from each concern, controlling for segment. Total engagement was operationalized as the combination of all engagement scores (range: 5 to 20) and each concern was coded as a binary variable where 1 is ‘extremely’ or ‘very concerned’. We also looked at interaction between concern and segment to determine if segment moderated the effect of the concern on willingness to engage.

## Acknowledgements

Funding: The authors acknowledge that they received no funding in support for this research.

## Author contributions

Each author’s contribution(s) to the paper should be listed (we suggest following the CRediT model with each CRediT role given its own line. No punctuation in the initials.

Examples:

Conceptualization: KH, TS, ATA

Methodology: CM, SM, KB

Investigation: ATA, CM, KH

Visualization: CM

Supervision: SJE, JLS, EH

Writing—original draft: SBB, DLA

Writing—review & editing: SBB, DLA, PRB, EH

## Data and materials availability

All data are available upon request.

