## Supplemental Material for "Public engagement with biomedical science: An analysis of Black, Hispanic, and General Population adults in the United States"

Adina T. Abeles *et al.*

**This PDF file includes:**

Supplementary Text
Tables S1 to S2

### Supplementary Text

Survey details

We conducted three national probability sample surveys of adults age 18 and older in each of the three populations of interest (Black (N= 963), Hispanic (N=964), General population (N=957)) using the NORC at the University of Chicago's AmeriSpeak panel.

Funded and operated by NORC at the University of Chicago, **AmeriSpeak®** is a probability-based panel designed to be representative of the US household population. Randomly selected US households are sampled using area probability and address-based sampling, with a known, non-zero probability of selection from the NORC National Sample Frame. These sampled households are then contacted by US mail, telephone, and field interviewers (face to face). The panel provides sample coverage of approximately 97% of the U.S. household population. Those excluded from the sample include people with P.O. Box only addresses, some addresses not listed in the USPS Delivery Sequence File, and some newly constructed dwellings. While most AmeriSpeak households participate in surveys by web, non-internet households can participate in AmeriSpeak surveys by telephone. Households without conventional internet access but having web access via smartphones are allowed to participate in AmeriSpeak surveys by web. AmeriSpeak panelists participate in NORC studies or studies conducted by NORC on behalf of governmental agencies, academic researchers, and media and commercial organizations.

Panelists were offered the cash equivalent of $3 for completing the survey. The survey was fielded from October 21 to November 6, 2021. Respondents classified as rushers (those who completed the survey in less than ⅓ of the median completion time), straightliners (respondents who straightlined responses on three or more segmentation battery grids), or skippers (respondents who skipped more than 50% of total questions) were removed from the sample (N = 126).

Panelists are recruited to AmeriSpeak using area probability and address-based sampling; sample households are contacted by U.S. mail, telephone, and field interviewers. A panel base sampling weight was applied to account for probability of selection into the panel, to align with U.S. Census benchmarks for each population on age (18-24, 25-29, 30-39, 50-59, 60-64, 65+), gender (male and female), education (Less than high school, high school/GED, Some college, BA and above), race/ethnicity (NonHispanic White, Non-Hipsnic Blakc, Hispanic, Non-Hispanic Other), housing tenure (Home owner and other), telephone status (cell only, landline only/phoneless, both cell and landline), and Census Division (new England, Middle Atlantic, East North Central, West North Central, South Atlantic, East South Central, West South Central, Mountain, Pacific), age x gender (18-34 Male, 18-34 female, 35-49 male, 35-40 female, 50-64 male, 50-64 female, 65+ male, 65+ female), and age x race/ethnicity (18-34 non-Hispanic White, 18-34 all other, 35-49 non-Hispanic White, 35-49 all other, 50-64 non-Hispanic White, 50-64 all other, 65+ Non-Hispanic White, 65+ all other), and to account for potential nonresponse bias among AmeriSpeak panelists. Post-stratficiation weights were also applied to account for any deviations from the US Census benchmarks for each population.

43 Segmentation Questions

Dimension: Curiosity and open-minded thinking

- I prefer news that aligns with or supports my beliefs.
- I consult multiple sources to make difficult decisions.
- The information I get from Google is sufficient to make most of my difficult decisions.
- I update my beliefs in response to new information or evidence.
- I am usually not convinced by an opposing argument.
- It's important to me that I'm usually right.
- Allowing oneself to be convinced by an opposing argument is a sign of good character.
- As I grow older, I've become less interested about the world around me.
- I'm comfortable publicly admitting when I'm wrong.

Dimension: Hierarchical/egalitarian

- The government interferes far too much in our everyday lives
- It's not the government's business to try to protect people from themselves
- The government should do more to advance society's goals, even if it means limiting the freedom and choices of individuals
- We have gone too far in pushing equal rights in this country
- Our society would be better off if the distribution of wealth was more equal
- Respect for authority is something all children need to learn
- Men and women each have different roles to play in society

Dimension: Generalized Trust

- Generally speaking, I trust other people.
- Generally speaking, other people treat me unfairly.
- Generally speaking, other people are helpful to me.

Dimension: Scientific thinking

- I follow a scientific approach when answering questions
- I trust my gut more than experts when making important decisions.
- Personal experience is more valuable than facts and statistics when making decisions.
- The science education I received in school helps me in my everyday life.
- I don't feel qualified to have a conversation about scientific research.

Dimension: Science in Society

- My local school board should make more decisions based on scientific thinking
- Scientists don't think about how their work affects the average person
- Scientists are arrogant
- Science is just another tool of manipulation that those in power can use to push their agenda.
- My city council should make more decisions based on scientific thinking.
- I'd pay more taxes to fund scientific research that directly benefits my local community.
- I'd pay more taxes to fund science education in my local school district.
- Most scientific research is driven by special interests that don't care about the average person.

Dimension: Role of biomedical science in society

- Medical science creates just as many problems as it solves
- Medical scientists should take an active role in debates about public health issues
- The decisions of my local health authority should be based on medical science more than economics
- Most medical research is driven by special interests that don't care about people like me
- The work of medical science researchers helps my community
- Progress in medical science research should be a top priority for elected officials

Dimension: Deference to biomedical authority

- Medical doctors have my best interest in mind when they suggest remedies
- I trust medical researchers to make ethical decisions
- It is important for medical scientists to get research done even if they displease people by doing it
- Medical scientists should make the decisions about which types of medical research will help society.
- Medical doctors have our best interest in mind when they tell the public what to do.

<u>Additional variables to describe the segments</u>
The following questions were included to help describe the segments after the cluster analysis. These were not included in the non-negative matrix factorization model.

*Willingness to participate in medical research*
Participants were asked, "How likely are you to do any of the following activities, if the opportunity arises?" And were shown the following activities in random order:

- Participate in medical research at a local university
- Participate in a vaccine trial at a local university
- Donate tissue to a local university for medical research
- Participate in medical research for the federal Centers for Disease Control
- Donated money to support medical research

The response options were "Extremely likely, very likely, somewhat likely, not at all likely".

*Concerns with engagement*
Participants were asked, "There are lots of reasons some people do NOT want to participate in medical research studies. When you think about participating in medical research studies, how concerned are you about each of the following?" And were shown the following concerns in random order:

- Getting COVID-19
- Getting a disease other than COVID-19
- Being a 'guinea pig'
- Results not being confidential
- Having to pay for research treatments
- The trustworthiness of the research process or the researcher

The response options were "Extremely concerned, very concerned, moderately concerned, slightly concerned, not at all concerned".

*Engagement with biomedical sciences*
Participants were asked, "Thinking back over the last 6 months, about how often have you done the following activities?" And were shown the following activities in random order:

- Read about medical issues in books, newspapers, magazines, scientific journals, etc.
- Talk to family, friends, and other acquaintances about medical related issues
- Discuss or share medical or health related issues on social media
- Read about health and medical-related issues on social media
- Encourage other members of your community to get a vaccine

The response options were, "Very often, Somewhat often, Rarely, Never"

*Trust*
Participants were asked, "How much do you trust each of the following to do what is right for you or your family?" And were shown the following in random order:

- University medical research center
- Primary health care provider (e.g., local health care clinic, primary care office)
- Local public health agency
- The federal government (Centers for Disease Control, National Institute of Health, Federal Drug Administration)
- A pharmaceutical company

*Trust in medical information source*
Participants were asked, "To what extent do you trust the medical information that you gather from the following sources?" And were provided the following in random order:

- TV
- Radio
- Internet
- Social media
- Family
- Friends

And the response options were, "Trust it a great deal, Mostly trust it, mostly distrust it, strongly distrust it, Don't use as a source of information".

*Doctor visit regularity*
About how long has it been since your last regular check up with a doctor?

1. Less than six months ago
2. Between six months to 1 year
3. 1-2 years
4. More than 2 years

*Trusted doctor*
When you have a concern about your health or the health of an immediate family member, do you have a medical professional that you trust to talk to?

1. Yes, more than one
2. Yes, one
3. No, not even one

*Perceived discrimination*
How often, if ever, do you believe people in your community experience discrimination or are treated poorly when seeking medical care because of their race, ethnicity, or the color of their skin?
1. All the time
2. Most of the time
3. About half the time
4. Some of the time
5. Never

*Lived discrimination*
What about you? How often, if ever, have you or anyone in your immediate family experience discrimination or been treated poorly when seeking medical care due to your race, ethnicity or color of your skin?
1. All the time
2. Most of the time
3. About half the time
4. Some of the time
5. Never

*Avoid treatment*
How often, if ever, have you avoided seeking medical treatment for you or others in your immediate family out of concern that you would be discriminated against or treated poorly because of your race, ethnicity or color of your skin?
1. All the time
2. Most of the time
3. About half the time
4. Some of the time
5. Never

*Community action*
Which, if any, of the following activities have you completed in the previous 12 months?
1. Contacted the editor of a local local newspaper or magazine
2. Contacted a local elected official about a particular issue
3. Organized an event to help improve my community
4. Volunteered my time to an important local cause or organization
5. Donated money to an important local cause
6. Served on a committee for a local club or organization
7. Written online about an important local cause
8. Have not taken any of the above actions

*News - general*
Where do you get most of your news from?
1. Television
2. News websites or apps
3. Through social networking sites (such as Facebook or Twitter)

4. The radio
5. Podcasts
6. In print
7. Other (specify): ___________ [ANCHOR]

*News - specific*

Which news outlet do you turn to for news? Select up to three.

1. ABC
2. CBS
3. CNN
4. Fox News
5. MSNBC
6. NBC
7. PBS
8. Local TV
9. Breitbart
10. Buzzfeed
11. Google News
12. The Huffington Post
13. NPR
14. Yahoo News
15. The New York Times
16. The Washington Post
17. The Wall Street Journal
18. USA Today
19. Univision
20. Radio One/Media One
21. Local Newspaper
22. Other (specify): ___________
23. None of the above

Additional details on methods

The objective of NMF algorithms is to solve problems iteratively, by building a sequence of matrices that reduces at each step the value of the objective function. In the simplest form, NMF consists in finding an approximation of

$$X \approx W * H$$

Where X is a non-negative matrix of *n(respondents)*p(variables);*

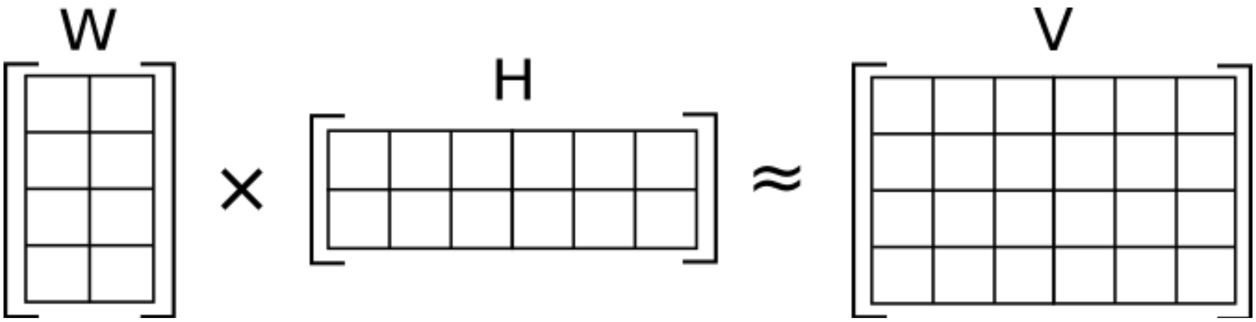


NMF can further be specified as:

$$\min_{W,H\geq 0} \underbrace{[D(X,WH)+R(W,H)]}_{=F(W,H)}$$

Where: $D$ is a loss function that measures the quality of the approximation; $R$ is an optional regularization function, defined to enforce desirable properties on matrices $W$ and $H$; and $F$ is the objective function ($W_k$ and $H_k$ reduce at each step of $F$).

Ultimately, the NMF works to cluster like-minded respondents into segments by using the segmentation battery items.

Like any other clustering algorithm, we can judge the performance of the model by checking the dispersion, silhouette and sparseness scores, along with a generic size breakdown for the given *K*'s.

- Dispersion Score: measures the reproducibility of the clusters obtained from NMF, ranges between 0 and 1, with 1 being the highest possible score
- Silhouette Score: measures for each observation how properly it fits the segment (1) in comparison to the other closest segment (-1), final score is the mean for all observations
- Sparseness score: measures how much energy of a vector (or matrix) is packed into only a few components. Sparseness is a real number ranging from 0 to 1. It is equal to 1 if and only if x contains a single nonzero component, and is equal to 0 if and only if all components of x are equal. It interpolates smoothly between these two extreme values. The closer to 1 is the sparseness the sparser is the vector.

Relative segment size is also important in selecting a segmentation solution. Ideally, segments are not so big such that they lack any real differentiating power (e.g. greater than 50% of the population) or so small such that they represent only a niche portion of the population (e.g. less than 5%). It is also useful to note that there is not necessarily one obvious solution, and that multiple characteristics of each solution are evaluated against each other in order to select the best fit. For example, size may be balanced against silhouette scores or the qualitatively-defined consistency of a mindset.

**Table S1: Breakdown of Possible Segmentation Solutions by Population**

| | Segmentation Solution | Segment 1 Size | Segment 2 Size | Segment 3 Size | Segment 4 Size | Segment 5 Size | Segment 6 Size |
|---|---|---|---|---|---|---|---|
| Gen. Pop. | 4-Segment Solution | 4% | 17% | 31% | 48% | – | – |
| | 5-Segment Solution | 19% | 32% | 13% | 27% | 9% | – |
| | 6-Segment Solution | 4% | 18% | 19% | 21% | 21% | 17% |
| Black Pop. | 4-Segment Solution | 28% | 11% | 36% | 25% | – | – |
| | 5-Segment Solution | 13% | 28% | 10% | 34% | 16% | – |
| | 6-Segment Solution | 40% | 7% | 7% | 23% | 20% | 3% |
| Hisp. Pop. | 4-Segment Solution | 26% | 24% | 29% | 21% | – | – |
| | 5-Segment Solution | 9% | 24% | 29% | 19% | 19% | – |
| | 6-Segment Solution | 22% | 3% | 7% | 14% | 23% | 31% |

*Synthesizing the segments*

NMF allows us to predict the cluster assignment, which is computed as the index of the dominant basis component for each respondent (e.g., the highest probability the respondent would fit in any of the clusters). Using the cluster membership, we can start describing our segments by understanding how they answered the segmentation battery questions.

To do that, we create index values for each of the questions for each segment. We define an index value as the percentage of respondents in a segment that agreed ("Strongly agree" and "Somewhat agree" on the Likert scale) with the statement divided by the overall agreement of all respondents times 100.

$$Index = \frac{\%\ \mathrm{agree}_{\mathrm{segment}}}{\%\ \mathrm{agree}_{\mathrm{population}}} * 100$$

An index value of 100 means that the segment agreement doesn't differentiate from the overall population. Index values above 100 indicate that the segment agrees more with the statement compared to the overall population, and an index value below 100 indicates that the segment agrees less with the statement.

Given that each segmentation battery item belongs to one of the seven mindset categories, the high- and low-indexing items correspond to high- or low-propensity associations with the categories. For example, the *Segment 5* from the general population is defined by hierarchical thinking, but open-mindedness does not play a large role in understanding that segment's prevailing mindset toward science. This can be observed descriptively: The items with the *highest* index values are "Science is just another tool of manipulation that those in power can use to push their agenda" and "We have gone too far in pushing equal rights in this country." The latter statement is one of the seven items on the hierarchical thinking battery, and other statements from that battery (e.g. "It's not the government's business to try to protect people from themselves" or "The government interferes too much in our everyday lives") also have relatively high index values. Just as important, and adding credibility to the *Segment 5* characterization, the lowest and third-lowest index values are ones that would have indicated a more communitarian/egalitarian mindset, which also provides strong evidence of hierarchical thinking among this group.

In assessing the segments for each of the three populations, it became clear that there is segment overlap between the populations: *Segment 1* appears in all three populations, *Segment 3* and *Segment 5* exist in both the general and Hispanic populations, and we find *Segment 4* in the Black and general populations. Thus, there are five segments per population and ten total unique segments.

Beyond the qualitative assessment of the segments, we aggregated item indexes as a robustness check up to the category level and generated a standardized score for each category by segment. Segment category scores are presented in Table 2.

**Table S2: Standardized Segment Propensity Scores by Segmentation Category**

| Segment | Category | | | | | |
|---|---|---|---|---|---|---|
| | **Scientific Thinking** | **Open-Mindedness** | **Generalized Trust** | **Science in Society** | **Hierarchical Thinking** | **Deference to Biomedical Authority** |
| Segment 1 | 4 | 4.5 | 3.5 | 5 | 0.5 | 4 |
| Segment 2 | 5 | 4.5 | 5 | 4.5 | 3.5 | 4.5 |
| Segment 3 | 0.5 | 1 | 1 | 1 | 2.5 | 2 |
| Segment 4 | 2 | 3 | 0.5 | 0.5 | 2 | 0.5 |
| Segment 5 | 2 | 3.5 | 4.5 | 0.5 | 5 | 0.5 |
| Segment 6 | 2 | 5 | 0.5 | 2.5 | 1 | 0.5 |
| Segment 7 | 4 | 4 | 4.5 | 4.5 | 1.5 | 5 |
| Segment 8 | 0.5 | 1.5 | 0.5 | 1 | 2 | 3.5 |
| Segment 9 | 1 | 1.5 | 2.5 | 0.5 | 3 | 1 |
| Segment 10 | 1 | 0.5 | 0.5 | 1 | 2 | 3.5 |

The standardization aligns well with our qualitative segment assessments. The approach to standardizing the segment category scores is as follows:

1. Classify each segmentation item into the correct segmentation category.

2. Create a column for reverse-coded values. All items should be coded such that a *high* index value on that item also means a *high* propensity within that mindset category, and vice versa. The column takes on a value of -1 if it *should* be reverse-coded and 1 if it should not.
3. An index value of 100 means that the item was not at all useful in differentiating the segments. Subtract 100 from the index value to get the item's distance from 100.
4. Take that value and multiply it by the reverse-coding column of 1s and -1s. Now each item has a corresponding value telling us how far each value is from 100 for each segment.
5. For each segment, calculate each *category*'s mean distance from 100. In other words, account for the total effect of a category's items on differentiating the segments.
6. Index those values on a 0 to 1 scale for each category and when comparing all segments, with 0 corresponding to the *lowest* value for each category and 1 corresponding to the *highest* value.
7. Proportionally allocate values from 0.5 to 5 (in increments of 0.5) based on the 0 to 1 scale values given to each segment for each category.

It should be noted that the values start at 0.5 to avoid the impression that any particular segment is completely "lost" or "hopeless" for a given segmentation category. Additionally, in this case a value of 5 is restricted to the segment that is *most* defined by a particular category, so it is easiest to differentiate the segments. The end-result of this process is ten unique segments, five total per population, that are rigorously differentiated yet comparable to one another on key attitudes toward science.
